# FcRn, but not FcγRs, drives maternal-fetal transplacental transport of human IgG antibodies

**DOI:** 10.1101/2020.03.22.999243

**Authors:** Sara Borghi, Stylianos Bournazos, Natalie K. Thulin, Chao Li, Anna Gajewski, Robert Sherwood, Sheng Zheng, Eva Harris, Prasanna Jagannathan, Lai-Xi Wang, Jeffrey V. Ravetch, Taia T. Wang

**Affiliations:** The Laboratory of Molecular Genetics and Immunology, The Rockefeller University, 1230 York Avenue, New York, NY 10065, USA; Department of Medicine, Stanford University School of Medicine, Stanford, CA 94305, USA; Department of Chemistry and Biochemistry, University of Maryland, College Park, MD 20742, USA; Sustainable Sciences Institute, Managua, Nicaragua; Proteomics Facility, Institute of Biotechnology, Cornell University, Ithaca, NY, 14853, USA; University of California, Berkeley, School of Public Health, Berkeley, CA 94720, USA; Department of Microbiology and Immunology, Stanford University, Stanford, California 94305, USA; Chan Zuckerberg Biohub, San Francisco, CA 94518, USA

**Keywords:** Immunoglobulin, IgG, placental transfer, Fc domain, FcRn, Fcγ receptors

## Abstract

The IgG Fc domain has the capacity to interact with diverse types of receptors, including FcRn and FcγRs, which confer pleiotropic biological activities. Whereas FcRn regulates IgG epithelial transport and recycling, Fc effector activities, such as ADCC and phagocytosis are mediated by FcγRs, which upon crosslinking transduce signals that modulate the function of effector leukocytes. Despite the well-defined and non-overlapping functional properties of FcRn and FcγRs, recent studies have suggested that FcγRs mediate transplacental IgG transport, as certain Fc glycoforms were reported to be enriched in fetal circulation. To determine the contribution of FcγRs and FcRn to the maternal-fetal transport of IgG, we characterized the IgG Fc glycosylation in paired maternal-fetal samples from patient cohorts from Uganda and Nicaragua. No differences in IgG1 Fc glycan profiles and minimal differences in IgG2 Fc glycans were noted, whereas the presence or absence of galactose on the Fc glycan of IgG1 did not alter FcγRIIIA or FcRn binding, half-life, or their ability to deplete target cells in FcγR/FcRn humanized mice. Modeling maternal/fetal transport in FcγR/FcRn humanized mice confirmed that only FcRn contributed to transplacental transport of IgG; IgG selectively enhanced for FcRn binding resulted in enhanced accumulation of maternal antibody in the fetus. In contrast, enhancing FcγRIIIA binding did not result in enhanced maternal/fetal transport. These results argue against a role for FcγRs in IgG transplacental transport, suggesting Fc engineering of maternally administered antibody to only enhance FcRn binding as a means to improve maternal/fetal transport of IgG.

**Significance Statement:** Transport of IgG antibodies from the maternal to the fetal circulation is a key process for neonatal immunity, as neonates cannot sufficiently generate IgG antibodies to reach protective levels during the first months after birth. In humans and other primates, maternal to fetal transport of IgG antibodies is largely mediated through the placental tissue. FcRn has been previously identified as the major driver of IgG transplacental transport. Here we examined whether other receptors, such as FcγRs, also contribute to the maternal-fetal IgG transfer. By characterizing the Fc domain structure of paired maternal-fetal IgG samples and modeling transplacental IgG transport in genetically engineered mouse strains, we determined that FcRn, but not FcγRs, is the major receptor that mediates transplacental IgG transport.

## Introduction

Maternal to fetal transfer of IgG antibodies is central to neonatal immunity. This is because IgGs are not produced by the neonate at mature levels for nearly a year after birth, making maternal antibodies a principle mechanism of immunity during this vulnerable period of development. In species such as primates, this transfer is largely the result of transplacental transport of IgG. The neonatal Fc receptor (FcRn) is recognized as the primary transporter of IgGs across epithelial cells of the placenta. In addition to transport across placental cell layers, FcRn extends the half-life of IgGs through a mechanism of protective recycling that rescues IgGs from catabolic pathways. These functions of FcRn are possible because of high affinity interactions between FcRn and monomeric IgGs, triggered by acidification of endosomes after passive uptake of IgGs that are present in the extracellular space. The high affinity of FcRn-IgG interactions enables transport of IgGs within endosomes from the apical cell surface to the basolateral cell surface (in transplacental IgG transport) or back to the apical surface (for recycling of IgGs), where a return to neutral pH causes release of IgGs from FcRn back into circulation (1, 2).

FcRn interacts with the IgG heavy chain primarily at the CH3 domain of the IgG Fc. This is in contrast to interactions with Fc gamma receptors (FcγRs), which bind the hinge proximal region of the CH2 domain of IgG. Interactions between FcRn with IgGs are generally not thought to be impacted by IgG subclass or Fc glycosylation, while specific IgG allotypes (e.g. Gm3(b), Gm3(g)) may impact FcRn binding (3–6). This is in stark contrast to FcγR-IgG interactions which are profoundly impacted by both IgG subclass and Fc glycosylation (7, 8). While FcRn is recognized as the major transporter of IgGs from maternal to fetal circulation, important questions remain regarding the biology of transplacental IgG transport. These include the mechanism(s) governing preferential transfer of the IgG1 subclass and the variation of transfer efficiency that can occur based on IgG Fab specificity. Further, whether FcγRs have any role in transplacental IgG transport or whether specific post-translational modifications of IgGs can impact transfer through any mechanism are questions that have not been resolved (9–13).

An intriguing prospect for enhancing neonatal immunity has focused on the dynamics of transplacental transport and postulates that increasing transport of protective IgGs to the fetus during gestation could profoundly boost and extend neonatal immunity. This could be done by maternal administration of recombinant, anti-pathogen antibodies that are specifically engineered for increased placental transport. Through enhancing transfer of protective monoclonal antibodies, this strategy could reduce infant mortality while avoiding direct delivery of IgGs to newborns. Feasibility of this approach, however, requires that there is absolute clarity on mechanisms regulating the transport of IgGs across the placenta.

Recent studies suggested that post-translational modifications of IgGs and a specific low-affinity FcγR, FcγRIIIa, have a role in transplacental IgG transfer (14, 15). These studies characterized paired maternal and cord blood IgG, reporting a significant increase in digalactosylation on total cord IgGs (but not on antigen-specific IgGs) and suggesting that this modification may direct transplacental transfer of maternal IgGs in a process they term “placental sieving”. Through a series of binding and *in vitro* cellular assays, the authors conclude that digalactosylation of IgG increases the affinity of IgGs for both FcRn and FcγRIIIa and propose that FcγRIIIa contributes to transplacental transfer of galactosylated IgGs. Further, they report that digalactosylation enhances FcγRIIIa-mediated cellular activation.

In view of the importance of this topic and a prior study of IgG Fc glycosylation from paired maternal and cord IgGs that used rigorous methods for relative quantitation of Fc glycans and found no significant differences in IgGs transferred across the placenta (9), we have reexamined key conclusions from these studies. We characterized paired maternal and fetal IgGs from two separate clinical cohorts and have used a combination of *in vitro* approaches and *in vivo* functional studies to test the hypothesis that FcγRIIIa or other FcγRs may have a role in transplacental IgG transfer. In two clinical cohorts, we observed no consistent difference between digalactosylation of maternal and cord IgG1, the most abundant IgG subclass. Some small, but statistically significant differences were observed in IgG2 galactosylation between maternal and cord IgGs. IgG2 is the second most abundant IgG subclass and does not engage FcγRIIIa. Biochemical and *in vivo* functional studies designed to investigate whether galactosylation of IgG (either IgG1 or IgG2) impacts interactions with FcRn or FcγRIIIa found no evidence of modulation of IgG-FcγR or IgG-FcRn interactions through Fc galactosylation, *in vitro* or *in vivo.* To confirm the results, we developed a novel *in vivo* murine model in which human FcγRs and FcRn are expressed in place of their murine counterparts and found no evidence that enhancing IgG Fc binding to FcγRIIIa (or to other FcγRs) results in increased maternal to fetal IgG transport. In contrast, enhancing IgG-FcRn binding resulted in significantly increased transplacental transport of maternal IgGs. Our results thus support the conclusion that Fc glycosylation is unlikely to be a significant modifier of maternal to fetal IgG transport and that engineering IgGs to enhance FcγRIIIa binding will not result in enhanced transplacental transport.

## Results

### IgG Fc glycans do not predict the efficiency of transplacental IgG transfer

To investigate whether differences in posttranslational modifications, such as glycosylation, of IgGs are found in paired maternal/fetal samples and thus suggest a role in transplacental IgG transfer, we characterized the glycosylation of IgG Fc domains from paired maternal and venous cord (fetal) bloods from two, geographical distinct clinical cohorts (Table S1). One cohort of mothers and infants from which samples were studied is located in Tororo, a region of Uganda where malaria is endemic (16, 17). Some mothers in the cohort had a malaria infection during pregnancy while others remained free from malaria infection (Table S1A). From these Ugandan samples, we profiled both total and malaria specific, anti-circumsporozoite protein (CSP) IgGs. Samples from a second, Nicaraguan cohort were drawn during a period when Zika virus infections were endemic in the region. Mothers in the study had a PCR-confirmed ZIKV infection during pregnancy or had confirmed infection via the NS1 Blockade-of-Binding ELISA (BOB) assay (18) and had Zika envelope protein (E)-reactive IgGs in serum (Table S1B). From the Nicaraguan study, we profiled total IgGs and anti-Zika E IgGs.

To characterize Fc glycoforms, we employed a previously validated and sensitive method (19), in which trypsin-digested, purified IgGs are subjected to nanoscale liquid chromatography coupled to tandem mass spectrometry (nano LC-MS/MS), enabling relative quantification of IgG subclass-specific Fc glycoforms. Subclass-specific analysis of Fc glycans is critical for hypothesis generation around the potential function(s) associated with specific Fc modifications as the role of various Fc glycoforms on non-IgG1 subclasses is not yet defined. In samples from both cohorts, we observed no statistically significant differences between maternal and cord IgGs for the majority of Fc glycoforms (Figures 1 and 2, Table S2). In IgGs from the Ugandan cohort (Figure 1), there were no significant differences between maternal and cord Fc glycoforms of total or of anti-CSP IgG1. Analysis of IgG2 showed some small, but statistically significant differences in galactosylation of total IgG, with decreased monogalactosylation in cord blood, and increased digalactosylation and total galactosylation in cord, compared with maternal blood. Though statistically significant, each of these modifications varied by only ∼2% between maternal and cord IgGs (Table S2). In contrast, no differences were observed in anti-CSP IgG2. Total IgG (all subclasses) monogalactosylation was significantly different, with the mean abundance of monogalactosylation being ∼1.4% lower in cord, relative to maternal blood (Figure 1, Table S2). No differences in Fc glycosylation of anti-CSP IgGs (all subclasses) were observed (Figure 1). The largest mean difference observed in Fc glycans between cord and maternal blood from the Ugandan cohort was in monogalactosylation of bulk IgG2, with cord blood having ∼2.3% lower levels than maternal blood (Figure 1, Table S2).

**Figure 1.**
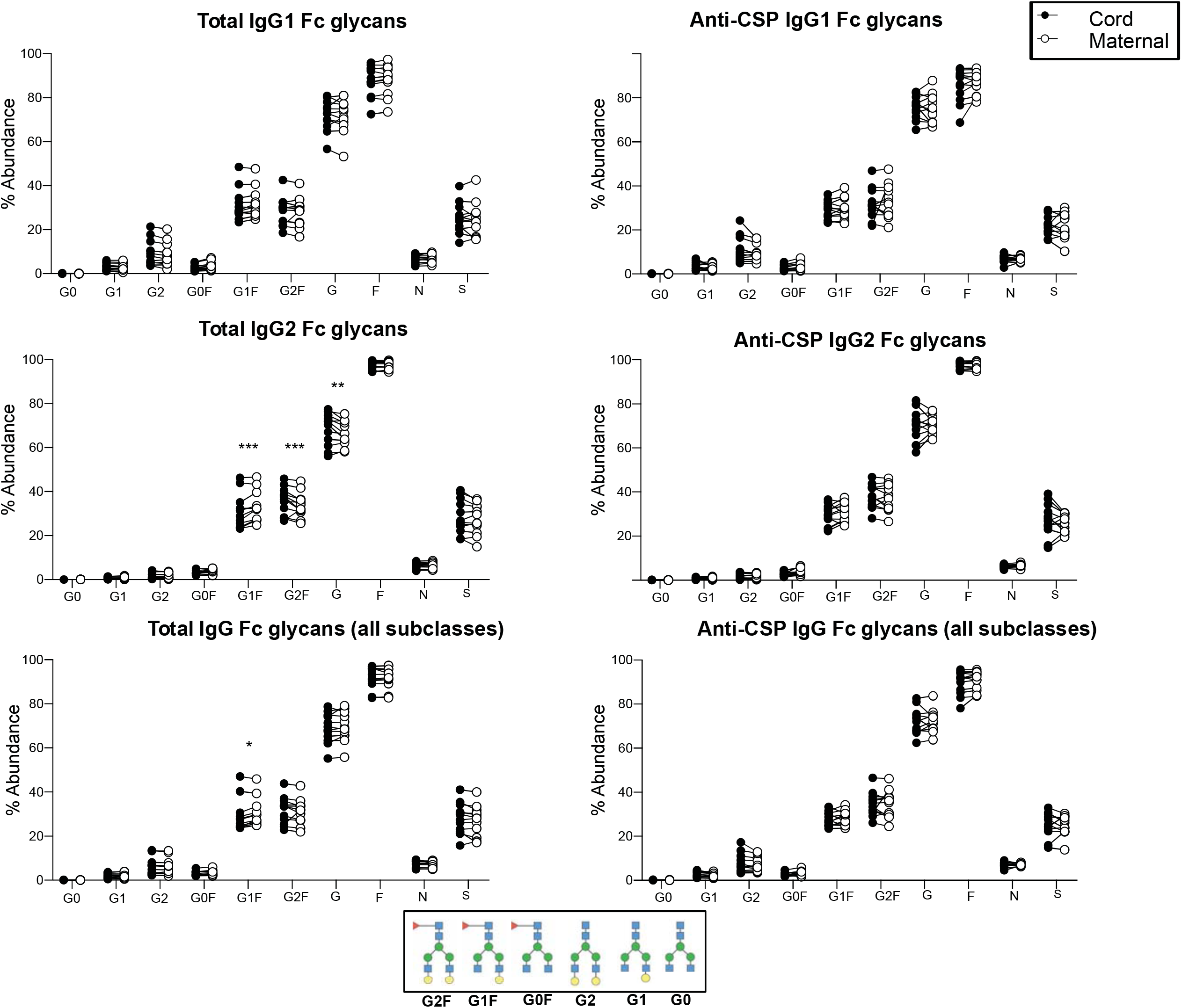
Relative abundance of IgG Fc glycans in paired maternal and cord bloods in the Ugandan cohort. Total or anti-malaria circumsporozoite protein (CSP) IgGs were purified from maternal or venous cord blood and the relative abundance of the following Fc glycoforms were characterized for each IgG category: G0, G1, G2, G0F, G1F, G2F, total galactosylation (G), total fucosylation (F), total bisection (N), total sialylation (S). Significance was assessed by 2-way ANOVA with Bonferroni’s multiple comparisons test where p<0.0125 was considered significant, *p < 0.0125, **p < 0.0025, ***p < 0.00025. Figure related to Tables S1 and S2.

In IgGs from the Nicaraguan study (Figure 2), total cord IgG1 levels of Fc monogalactosylation were reduced and were increased in sialylation relative to total maternal IgG1, while anti-E IgG1 total galactosylation was significantly elevated in cord blood (Figure 2). Total cord IgG2 was significantly elevated in total galactosylation and cord anti-E IgG2 digalactosylation and total galactosylation were elevated over maternal blood IgG (Figure 2). Total cord IgG (all subclasses) Fc monogalactosylation was significantly reduced, but was elevated in digalactosylated IgGs while cord anti-E IgG (all subclasses) was elevated in total galactosylation. The largest mean difference observed in Fc glycans between cord and maternal blood from the Nicaraguan cohort was in total galactosylation of bulk IgG2, with cord blood having ∼5.6% higher levels than maternal blood (Figure 2, Table S2).

**Figure 2.**
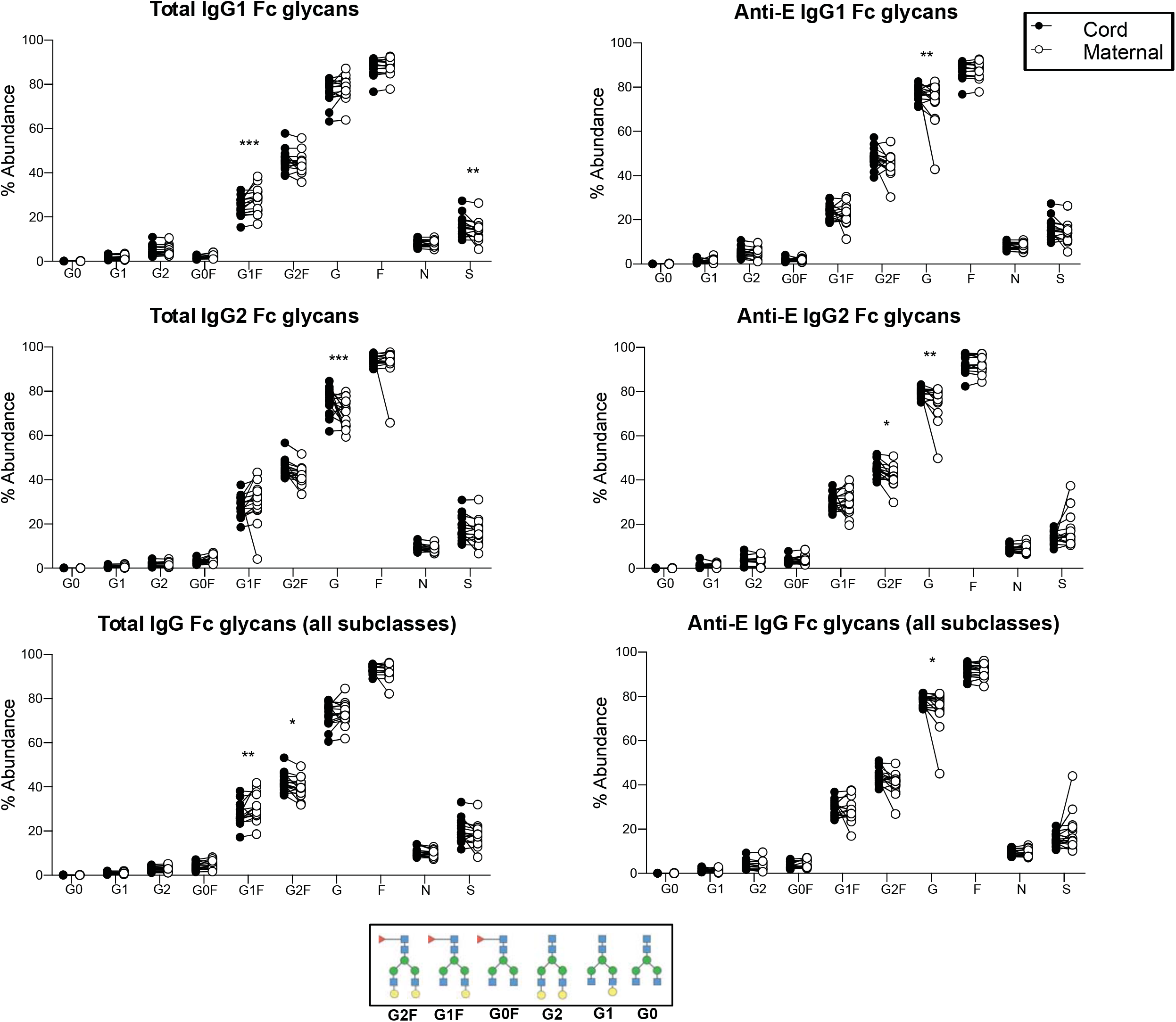
Relative abundance of IgG Fc glycans in paired maternal and cord bloods in the Nicaraguan cohort. Total or anti-Zika virus envelope (E) IgGs were purified from maternal or venous cord blood and the relative abundance of the following Fc glycoforms were characterized for each IgG category: G0, G1, G2, G0F, G1F, G2F, total galactosylation (G), total fucosylation (F), total bisection (N), total sialylation (S). Significance was assessed by 2-way ANOVA with Bonferroni’s multiple comparisons test where p<0.0125 was considered significant, *p < 0.0125, **p < 0.0025, ***p < 0.00025. Figure related to Tables S1 and S2.

Overall, we find that there were no consistent trends in Fc glycosylation between cord and maternal IgGs which would indicate a role for IgG glycosylation in transplacental transport of IgGs. Transplacental IgG transfer in these two clinical cohorts was not a function of Fc galactosylation (Figures 1,2, Table S2). Further, no difference between maternal and cord IgG fucosylation was observed, which could support a potential role for FcγRIIIa in transplacental IgG transport. With respect to IgG subclasses, results from the Ugandan and Nicaraguan cohorts were consistent with one another and with prior studies demonstrating a clear bias in transfer of IgG1 from maternal to cord blood and reduced transfer of the IgG2 subclass (Figure 3, Table S2) (20–22).

**Figure 3.**
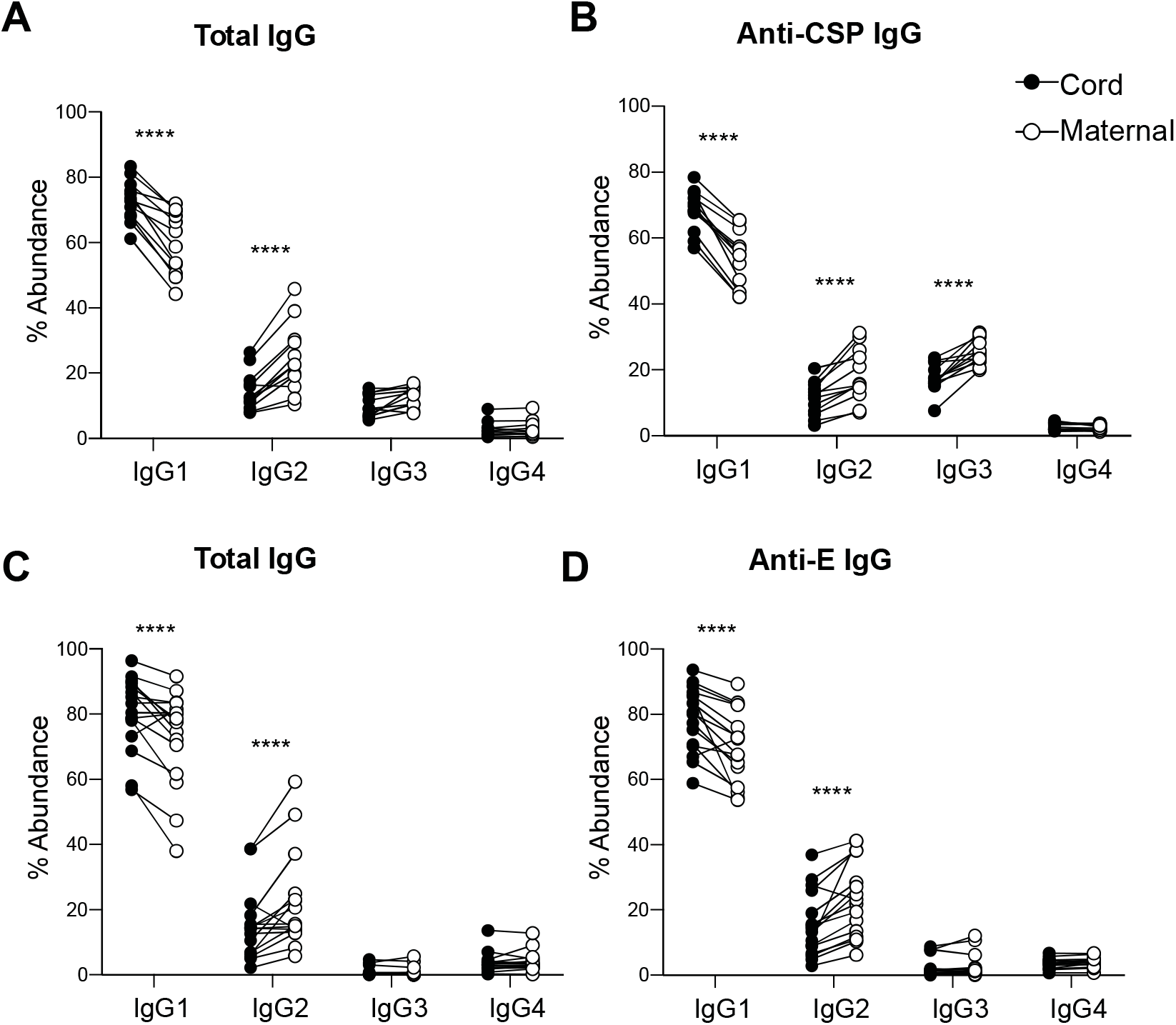
IgG subclass distribution in paired maternal and cord bloods. **(A,B)** Total or antimalaria circumsporozoite protein (CSP) IgG from the Ugandan cohort. **(C,D)** Total or anti-Zika virus envelope (E) IgG from the Nicaraguan cohort. Data from Figure 2 are quantified in Table S1. The abundance of specific IgG subclasses is shown. Significance was assessed by paired T-test with Bonferroni’s correction. \**P* < 0.05, \*\**P* < 0.01, \*\*\**P* < 0.001, \*\*\*\**P* < 0.0001. Figure related to Tables S1 and S2.

### Galactosylation of IgG Fc does not significantly impact binding to FcRn or FcγRs *in vitro*

Previous studies have established that core fucosylation of the Fc reduces affinity of IgG1 for FcγRIIIa, while galactosylation alone has no impact on FcγRIIIa binding and/or antibody-dependent cell-mediated cytotoxicity (23, 24). Galactosylation has been observed to increase Fc-FcγRIIIa binding, but only in absence of fucosylation (25). Recently, it has been reported that IgG galactosylation, in the presence of fucosylation, impacts IgG binding to FcγRIIIa and FcRn (14). Given these contrasting reports on the role of Fc galactosylation in modulating FcγR and FcRn affinity, we performed a comprehensive analysis of the FcγR and FcRn affinity of IgG Fc glycoforms with defined, homogenous glycan structures. Using a chemoenzymatic glycosylation remodeling method (26, 27), we generated homogenously agalactosylated (G0) or digalactosylated (G2) human IgG1 variants of the mAb 6A6 that specifically recognizes the murine platelet glycoprotein IIb (28). Production of G0 or G2 of anti-gpIIb mAb was further stratified by core fucosylation (G0F and G2F, respectively), and the homogeneity of the IgG glycoforms was assessed by LC-MS analysis (Figure S1).

Human IgG1 variants of 6A6 mAb were tested by surface plasmon resonance (SPR) for binding to all human FcγRs and to human FcRn. As expected, Fc fucosylation of mAb 6A6 significantly reduced binding to both FcγRIIIa^158F^ and FcγRIIIa^158V^ alleles (Figure 4A-B). In contrast, fucosylation did not significantly impact interactions with FcRn. The galactosylation status of the 6A6 mAb had no impact on FcγRIIIa (or other FcγR) or FcRn binding (Figure 4A-B, Figure S2A). To exclude mAb clone specific effects, the results of these SPR binding studies were confirmed using homogenous Fc glycoforms (G0 or G2) of the anti-influenza hemagglutinin (HA) mAb FI6 (Figure S3A).

**Figure 4:**
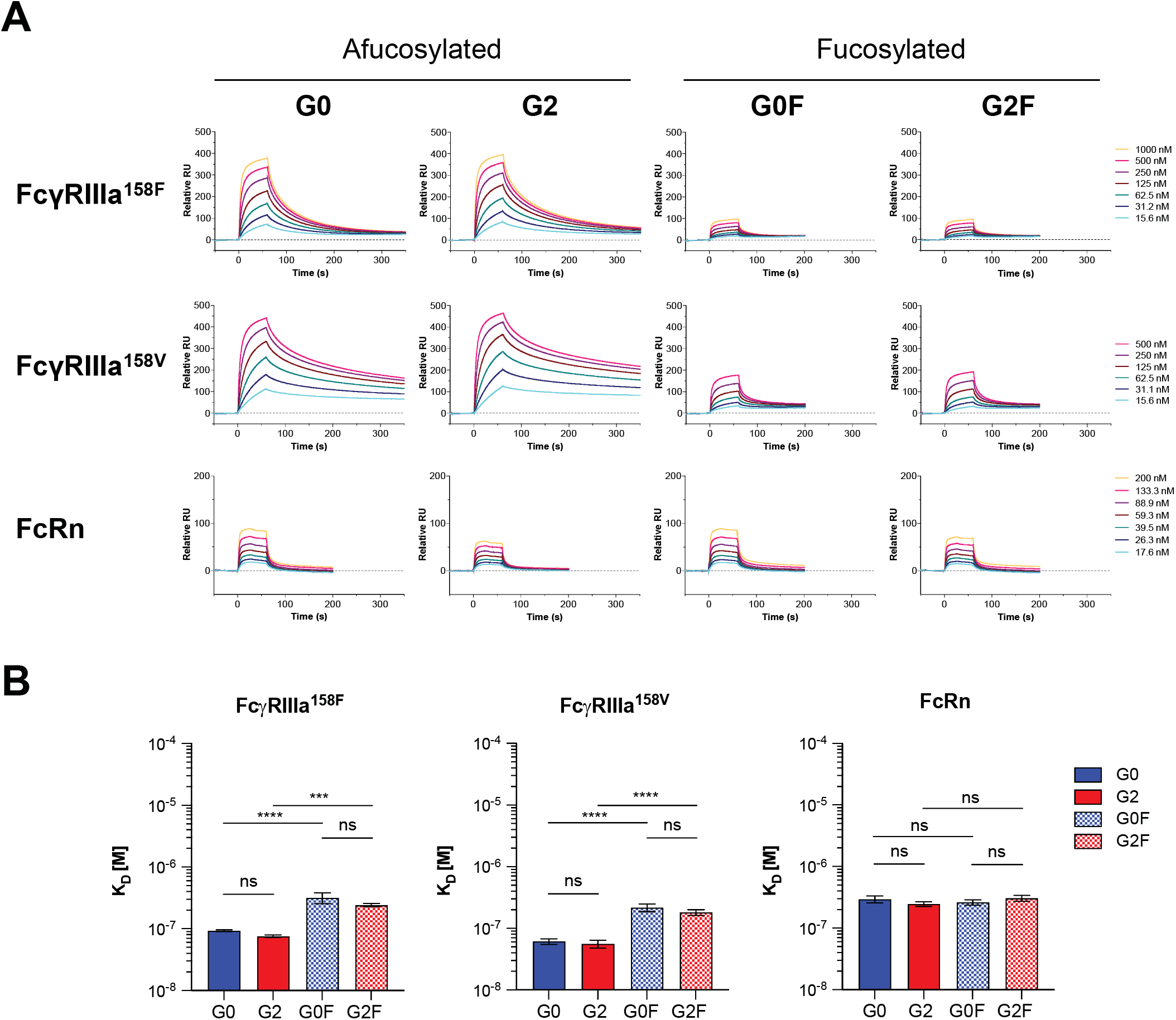
Effect of IgG1 core fucosylation and galactosylation on FcγRs engagement. (**A**) The affinity of glycoengineered mouse-human chimeric IgG1 of anti-gpIIb (6A6) mAb for human FcγRIIIa^158F^, FcγRIIIa^158V^ and FcRn ectodomains was determined by SPR analysis and representative SPR sensorgrams are presented. Data correspond to one experiment per interaction tested, which is representative of three independent experiments that gave similar results. (**B**) The equilibrium dissociation constant (K_D_, M) and the standard deviation are indicated for each tested interaction. Significance was assessed by one-way ANOVA followed by Bonferroni multiple comparison test. ns: not significant; ***p < 0.005; ****p < 0.0001. Figure related to Figures S1–S3.

Although analysis of the two clinical cohorts revealed no differences in Fc glycosylation between cord and maternal IgG1, a minor, yet statistically significant difference was evident in the levels of IgG2 Fc galactosylation levels between cord and maternal IgG (Figures 1,2 Table S2). To address whether Fc galactosylation of IgG2 impacts FcγR and/or FcRn affinity, we generated homogenously agalactosylated (G0) or digalactosylated (G2) human IgG2 variants of both anti-gpIIb 6A6 and anti-HA FI6 mAbs (Figure S1) and assessed their affinities for human FcγRs and FcRn by SPR. As observed for IgG1 (Figure 4), the galactosylation status of IgG2 had no impact on either FcγR or FcRn binding affinity (Figure 5A-B, Figure S2B). Thus, using homogeneously glycosylated IgGs revealed no significant differences in FcγR or FcRn binding affinities for galactosylated or agalactosylated Fc glycoforms for either the IgG1 or IgG2 subclasses.

**Figure 5:**
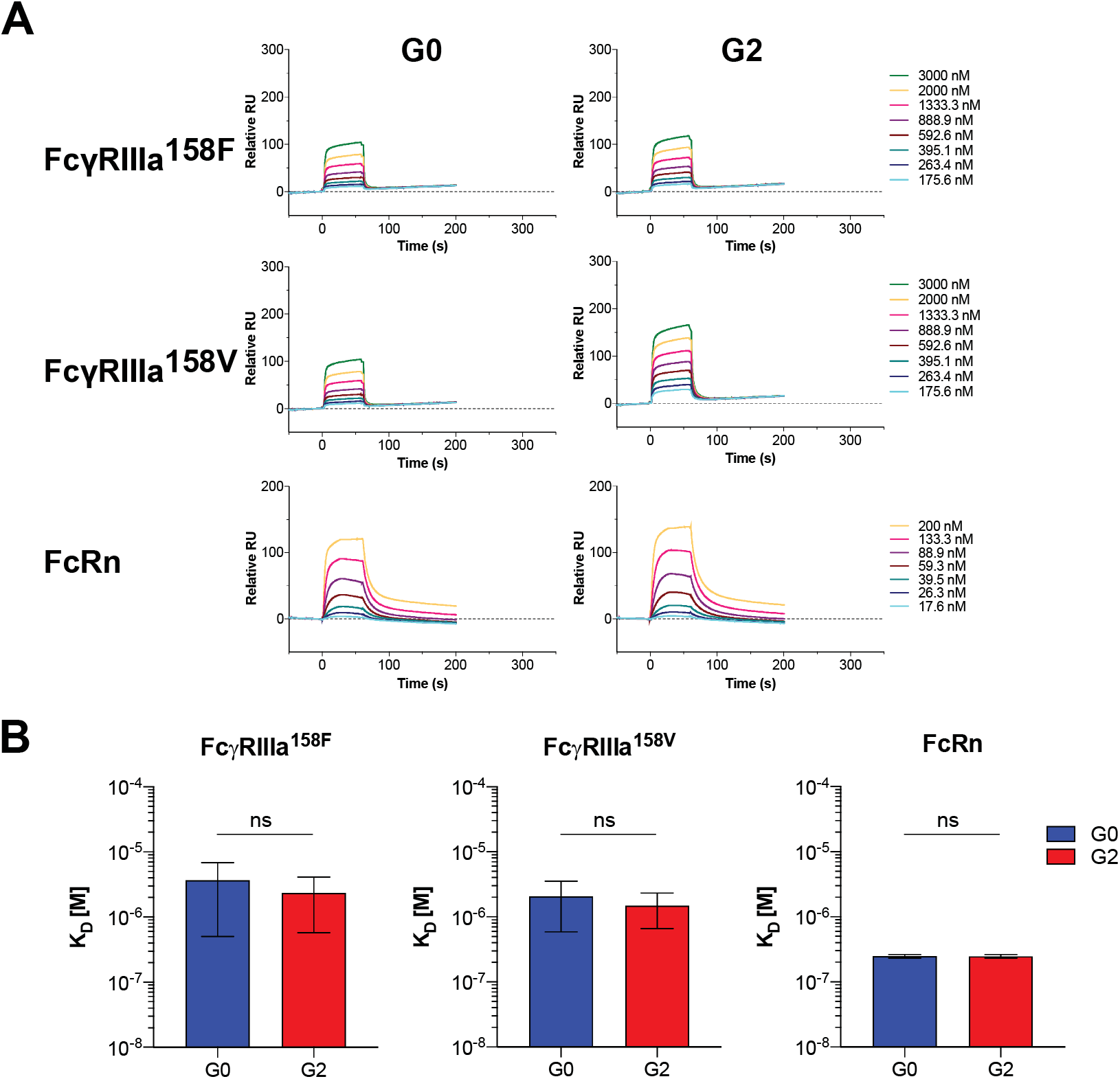
Effect of IgG2 core fucosylation and galactosylation on FcγRs engagement. (**A**) The affinity of glycoengineered mouse-human chimeric IgG2 of anti-gpIIb (6A6) mAb for human FcγRIIIa^158F^, FcγRIIIa^158V^ and FcRn ectodomains was determined by SPR analysis and representative SPR sensorgrams are presented. Data correspond to one experiment per interaction tested, which is representative of two or three independent experiments that gave similar results. (**B**) The equilibrium dissociation constant (K_D_, M) and the standard deviation are indicated for each tested interaction. Significance was assessed by unpaired t-test; ns: not significant. Figure related to Figures S1–S3.

### Galactosylation of IgG Fc does not significantly impact *in vivo* activity

To validate the physiological significance of these *in vitro* binding results, we assessed the *in vivo* effector function of Fc glycoforms of cytotoxic antibodies in FcγR humanized mice, which recapitulate the unique features of human FcγR physiology (29). To characterize the Fc effector activity conferred by Fc galactosylation, we took advantage of the well-defined *in vivo* platelet depleting activity of anti-gpIIb mAb 6A6. In mice that are fully humanized for FcγRs, administration of the chimeric mAb 6A6/hIgG Fc causes platelet counts to drop through the engagement of the activating FcγR, FcγRIIIa on myeloid cells. Platelet depletion is quantitatively increased as the binding affinity of the 6A6 human IgG Fc for FcγRIIIa is increased (29, 30). Thus, this system can be used to dissect how Fc modifications directly impact binding to FcγRIIIa and thereby its function. Using the homogenous glycovariants of human IgG1 and IgG2 mAb 6A6 previously characterized for their *in vitro* FcγR binding affinity (Figures 4,5), we observed that the absence of Fc fucosylation, regardless of galactosylation status, resulted in enhanced platelet depletion by mAb 6A6 (Figure 6). This was consistent with the increased affinity of afucosylated 6A6 glycoforms for FcγRIIIa, when compared with fucosylated 6A6 variants (Figures 4A-B, 5). In contrast, there was no difference in the activity of mAbs based on G0 or G2 glycoforms (Figure 6A, upper panel). Similarly, no difference was observed in the activity of G0 or G2 glycoforms in IgG2 subclass-mediated depletion (Figure 6A, lower panel).

**Figure 6:**
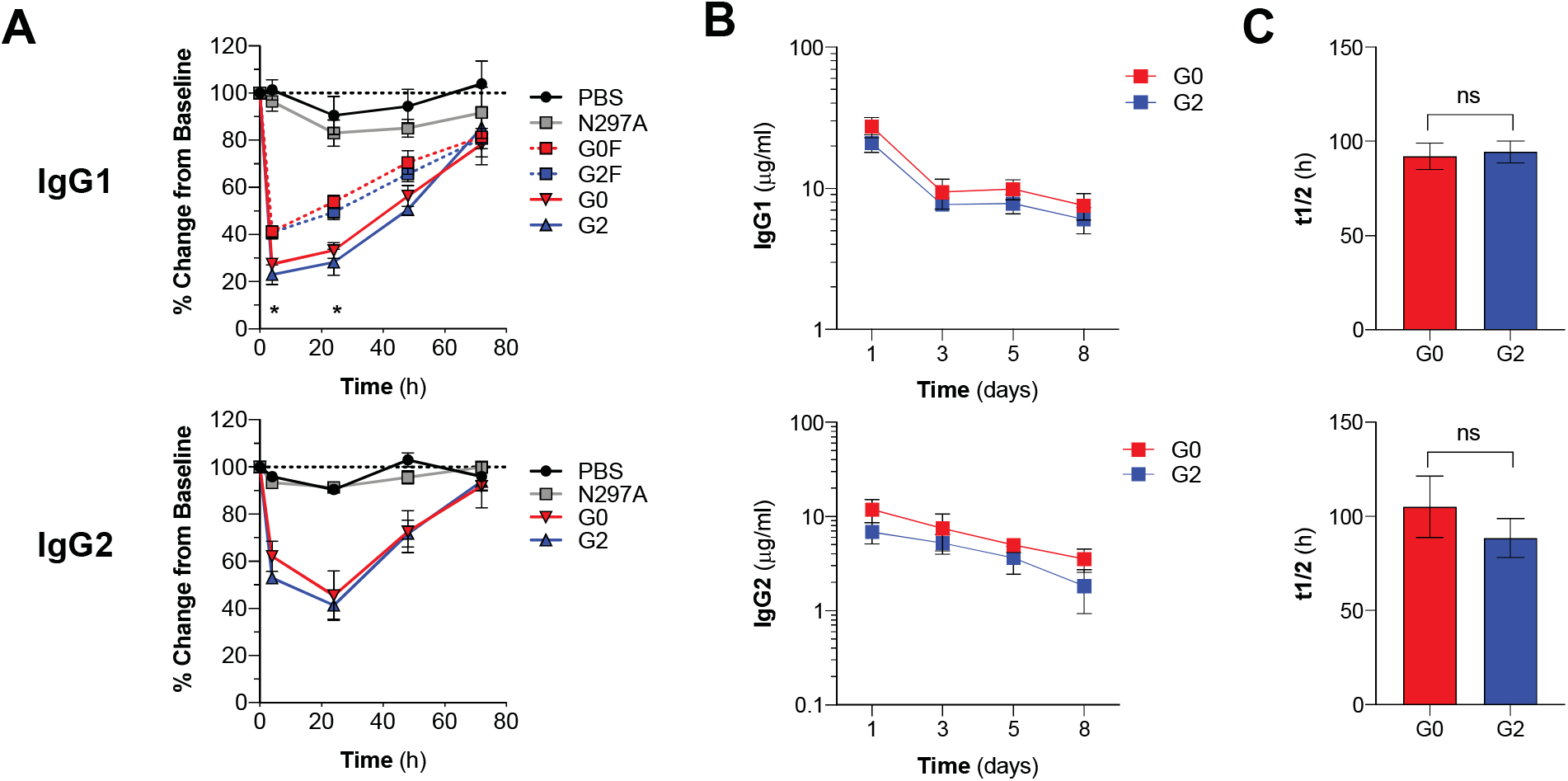
Evaluation of the *in vivo* Fc effector function and half-life of human IgG antibodies in FcγR and FcγR/FcRn humanized mice. (**A**) The cytotoxic activity of glycoengineered mAbs was evaluated in FcγR humanized mice. Timecourse analysis of platelet counts following administration of glycoengineered anti-gpIIb (6A6) IgG1 (upper panel) and IgG2 (lower panel) mAb (2 mg/kg) is presented as the mean ± SEM from 3-8 mice/group. The Fc domain variant N297A, lacking Fc glycosylation and therefore the FcγR binding ability, is used as negative control. Significance was assessed by two-way ANOVA followed by Bonferroni multiple comparison test; *p<0.05 G0 vs G0F and G2 vs G2F. (**B-C**) The *in vivo* halflife of glycoengineered anti-hemagglutinin (FI6) IgG1 (upper panels) and IgG2 (lower panels) mAb (50 μg/mouse) was evaluated in FcRn/FcγR humanized mice. Results are presented as the mean ± SD from 4-5 mice/group. Significance was assessed by unpaired t-test. ns = not significant. Figure related to Figures S4 and S5.

In addition to evaluating the Fc effector function of Fc glycovariants, we determined the pharmacokinetics (PK) profile of these glycovariants *in vivo.* Given the substantial interspecies differences between humans and mice in the affinity for human IgG antibodies for FcRn, we generated a novel mouse strain engineered to express all classes of human FcγRs and FcRn in lieu of the mouse orthologues of these receptors. Importantly, FcγR expression on effector leukocytes in the FcγR/FcRn humanized mice was consistent with expression on human leukocyte populations (Figure S4). To characterize the *in vivo* activity of human IgG antibodies in this novel mouse strain, we performed a series of studies to evaluate human FcRn and FcγR-mediated functions. Engineering IgG for higher affinity towards human FcRn using M428L/N434S (LS) mutations enhanced IgG half-life in the FcγR/FcRn-humanized mice (Figure S5A)(31). Cytotoxic antibody effector function was characterized using models of CD4^+^ T cell depletion and the mAb 6A6-mediated model of platelet depletion. As expected, chimeric anti-CD4 or 6A6 human IgG1 mAbs, Fc-engineered for higher affinity for activating FcγRIIA and IIIA and reduced for FcγRIIB, using the G236A/A330L/I332E (GAALIE) Fc mutations, were significantly more efficient at mediating depletion of CD4+T cells and platelets, respectively, compared with wild-type human IgG1 mAb variants or non-FcγR binding mAb variants (G236R/L328R (GRLR) Fc mutations) (Figure S5B-D) (32). Thus, human antibody functions mediated by human FcRn and FcγRs in this novel mouse model were intact.

To evaluate the role of IgG galactosylation status in the modulation of the FcRn-mediated IgG recycling, we compared the *in vivo* half-life of agalactosylated (G0) and digalactosylated (G2) Fc glycoforms of the anti-HA mAb FI6. Consistent with the SPR affinity data (Figures 4 and 5), the presence or absence of galactose residues had no significant impact on the *in vivo* half-life of either IgG1 (Figure 6B and C, upper panels) or IgG2 (Figure 6B and C, lower panels) variants of mAb FI6. Overall, our studies in FcγR/FcRn humanized mouse strains revealed that the galactosylation status of IgG1 and IgG2 antibodies had no impact on their capacity to interact with FcγRs or FcRn, modulating either IgG effector functions or half-life, respectively.

### Engineering IgG Fcs for enhanced binding to FcvRs does not impact transplacental IgG transfer *in vivo*

While our data do not support the contention that substantive changes in glycan profiles between maternal and fetal compartments are seen in matched samples from different patient cohorts, we nevertheless sought to directly test whether engineering IgGs for enhanced binding to any FcγR(s) might impact transplacental IgG transfer *in vivo.* To experimentally evaluate this possibility, we developed a novel mouse model using FcγR/FcRn humanized mice described above. Human IgG1 was transferred to pregnant mice one day prior to delivery and the levels of human IgG1 were characterized in the neonates. Human IgG1 levels in neonatal mice remained relatively unchanged in the 8 days following delivery when mice were housed with their biological mothers (Figure 7A). Next, we sought to determine whether this antibody was acquired by pups via transplacental transfer or via GI tract-mediated transfer after nursing (milk-acquired IgGs). To study this, we compared transfer of human IgG to pups who were housed after delivery with their birth mothers, who had been administered human IgG1 one day prior to delivery, or with surrogate mothers who were not administered any antibody prior to delivery. This comparison revealed that pups that did not receive human IgG via GI tract-mediated transfer had a rapid decline in serum IgG levels (Figure 7B,C). Thus, we determined that the level of transplacentally transferred IgG could be ascertained in the pups on the day of delivery, prior to feeding, a time point comparable to obtaining cord blood upon delivery in our human cohorts. To validate the GI tract-mediated IgG transfer, we studied IgG levels in naïve pups that were housed with mothers who were administered human IgG prior to delivery. Indeed, serum IgG levels rose rapidly in the pups, demonstrating substantial GI tract-mediated IgG transfer (Figure 7D). Thus, we concluded that the day of delivery was ideal for studying transplacentally transferred IgG in this novel model system.

**Figure 7:**
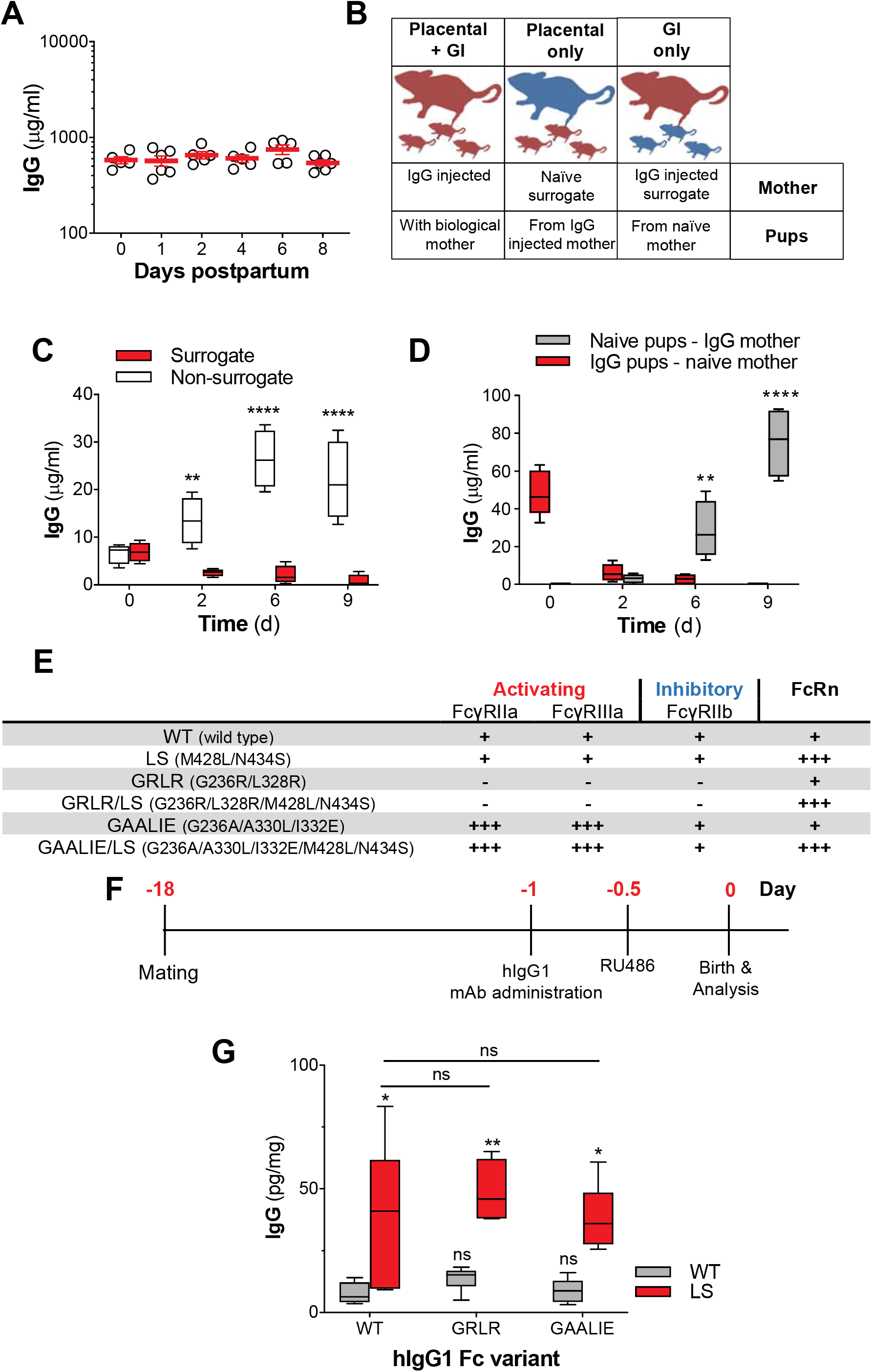
Evaluation of transplacental transfer of human IgG antibodies in FcRn/FcγR humanized mice. (**A**) Pregnant FcRn/FcγR humanized mice were injected with human IgG1 mAb (3BNC117) one day prior to delivery and the serum levels of human IgG1 were evaluated in the pups at different timepoint post-partum. (**B**) Experimental overview to differentiate placental-from GI tract-mediated maternal-to-fetal transfer of human IgG in FcRn/FcγR humanized mice involved the use of surrogate mothers that were injected or not with human IgG antibodies. (**C**) To assess the contribution of placental-mediated IgG transfer, pups born to mothers that were injected with human mAbs one day prior to delivery remained either with their biological mother (non-surrogate) or transferred to naïve surrogate mothers. Human IgG levels in the serum of pups were quantified at different timepoint following birth. n=4/group ****p<0.0001; **p=0.006 surrogate vs. non-surrogate. (**D**) Comparison of placental-(red boxes) and GI-tract-mediated (gray boxes) transfer of human IgG antibodies from maternal to fetal circulation. n=4-5mice/group; **p=0.001, ****p<0.0001 (**E**) FcγR and FcRn binding profile of Fc domain mutants with differential receptor affinity that were generated to study the contribution of human FcRn and FcγRs to the trans-placental transfer of human IgG antibodies. (**F**) Experimental strategy to evaluate the maternal-to-fetal transport efficacy of human IgG antibodies in FcRn/FcγR humanized mice. (**G**) Comparison of the human IgG levels in fetuses born to mothers previously injected with Fc domain variants of a human IgG1 mAb (3BNC117) n=3-9/group; ns= not significant vs WT; *p<0.02 vs WT or GAALIE; **p=0.003 vs GRLR. Figure related to Figures S4 and S5.

To determine whether FcγRs may have a role in transplacental IgG transfer, we developed variants of a human IgG1 mAb that were engineered for enhanced or reduced binding to specific human FcγRs or to human FcRn (Figure 7E). Human mAb variants were administered to pregnant mice (on day 19 of pregnancy) one day prior to induced delivery and antibody levels were quantified in pups upon delivery (Figure 7F). Engineering of mAbs for enhanced binding to activating FcγRs (FcγRIIa and FcγRIIIa), using the GAALIE mutations had no impact on transplacental IgG transfer (Figure 7G). Additionally, abrogating FcγR binding using GRLR mutations had no impact on transplacental IgG transfer. In contrast, engineering the Fc for enhanced affinity to human FcRn using LS mutations significantly increased the efficiency of transplacental IgG transfer (Figure 7G).

In summary, studies using this novel model of transplacental transfer of IgG demonstrated that, while FcRn could be harnessed to increase transplacental IgG transfer, modulating antibodies for their binding to FcγRs did not impact the transplacental transfer efficacy of human IgGs.

## Discussion

Here, using a variety of *in vitro* and *in vivo* approaches, we addressed questions also asked by recent studies as to whether FcγRs may be involved in transplacental transfer of human IgGs (9, 14, 15). Unlike some studies (14, 15) but consistent with other (9), we find that there is no enrichment for specific Fc glycoforms in cord blood over paired maternal blood. A significant difference in methodologies exists between these reports in that the Fc glycan analysis was not done in an IgG subclass-specific manner in these studies (14, 15); rather these studies characterized total IgG Fc glycans (antigen-specific IgG, or bulk IgG). This methodology does not account for the different activities that can be conferred by modification of different IgG subclasses by the same N-glycan. In contrast, the present study report relative quantification of Fc glycoforms on specific IgG subclasses. Together, our studies show no consistent trends among cohorts, indicating that Fc glycosylation is likely not a significant modifier of maternal-to-fetal IgG transport.

Our studies also find that Fc galactosylation had no statistically significant impact on FcγR binding, including FcγRIIIa, or on human FcRn binding for either the IgG1 or IgG2 subclasses. *In vivo* analysis of different IgG glycoforms in mice expressing human FcγRs and FcRn and lacking in their murine homologues confirmed that galactosylation had no statistically significant impact on FcγRIIIA-mediated effector function or FcRn-mediated half-life. Finally, direct analysis of transplacental transport of human IgG in a novel murine model demonstrated that modulating binding of maternal antibodies to FcγRs, including FcγRIIIA, had no impact on neonatal antibody concentration. In contrast, modulating maternal antibody FcRn affinity resulted in statistically significant increases in neonatal antibody concentration. While Jennewein et al. (14) propose that the enhanced FcγRIIIa-mediated effector function of cord blood IgG relative to maternal IgG was a function of enriched galactosylation of IgG in cord blood, we suggest that this activity is likely a consequence of the well-documented bias in transplacental transfer of the IgG1 subclass. IgG1 antibodies have enhanced affinity for activating FcγRs, including FcγRIIIa; thus, this is a straightforward hypothesis, explaining the difference in functional activity between maternal and fetal IgGs that was observed in their study.

We conclude that engineering antibodies for enhanced FcγRIIIa binding (or for binding to any FcγR) will not result in increased transplacental IgG transfer and is not a viable strategy to pursue for extending neonatal immunity. In contrast, our studies support the engineering of mAbs for enhanced FcRn affinity to increase the efficiency of IgG transfer from mother to fetus.

## Materials and Methods

### Clinical cohorts and samples

Paired maternal and cord blood samples were obtained from women and infants enrolled in PROMOTE (NCT02163447), a randomized clinical trial of novel antimalarial chemoprevention regimens in Eastern Uganda (16, 17). The study was approved by the Institutional Review Boards of the Makerere University School of Biomedical Sciences, the Uganda National Council for Science and Technology, and the University of California San Francisco. Written informed consent was obtained from all study participants.

Paired maternal and cord blood samples were obtained from the Nicaraguan Zika Positives study (NZP, approved by UC Berkeley IRB, protocol # 2016-10-9265, and the Nicaraguan IRB, protocol # NIC-MINSA/CNDR CIRE-19/12/16-078).The NZP study enrolled pregnant women confirmed as rRT-PCR ZIKV+ by the Ministry of Health or with a history of Zika-like symptoms during their pregnancy. For those who displayed Zika-like symptoms but did not have a ZIKV+ rRT-PCR result, recent ZIKV infection was confirmed via the NS1 Blockade-of-Binding ELISA (BOB) assay (18).

### IgG Fc glycan and IgG subclass analysis

Methods for relative quantification of Fc glycoforms and IgG subclasses have been described (19, 33). Briefly, IgGs were purified from serum by protein G purification. Antigen-specific IgGs were isolated on NHS agarose resin (ThermoFisher; 26196) coupled to the relevant protein. Following tryptic digestion of purified IgGs, nanoLC-MS/MS analysis was performed on tryptic peptides containing the N279 glycan using an UltiMate3000 nanoLC (Dionex) coupled with a hybrid triple quadrupole linear ion trap mass spectrometer, the 4000 Q Trap (SCIEX). Data were acquired using Analyst 1.6.1 software (SCIEX) for precursor ion scan triggered information dependent acquisition (IDA) analysis for initial discovery-based identification. For quantitative analysis of the glycoforms at the N297 site across the three IgG subclasses (IgG1, IgG2 and IgG3/G4), multiple-reaction monitoring (MRM) analysis for selected target glycopeptides, was applied using the nanoLC-4000 Q Trap platform to the samples after trypsin digestion. The m/z of 4-charged ions for all different glycoforms of the core peptides from three different subclasses as Q1 and the fragment ion at m/z 366.1 as Q3 for each of transition pairs were used for MRM assays. A native IgG tryptic peptide (131-GTLVTVSSASTK-142) with transition pair of, m/z 575.9^+2^ to mz 780.4 (y8^+^) was used as a reference peptide for normalization purposes. The IgG subclass distribution was quantitatively determined by nano LC-MRM analysis of tryptic peptides following removal of glycans from purified IgGs with PNGase F. Here, the m/z value of fragment ions for monitoring transition pairs was always larger than that of their precursor ions being multi-charged to enhance the selectivity for unmodified targeted peptides and the reference peptide. All raw MRM data was processed using MultiQuant 2.1.1 (SCIEX). MRM peak areas were automatically integrated and manually inspected. In the event that automatic peak integration by MultiQuant failed, manual integration was performed using the MultiQuant software.

### Preparation of homogeneous glycoforms of monoclonal antibodies

The glycoforms of monoclonal antibodies 6A6 and FI6 was synthesized by the chemoenzymatic glycan remodeling method following the previously reported procedure (26).

### Surface plasmon resonance (SPR)

Surface Plasmon Resonance experiments were performed with a Biacore T200 SPR system (Biacore, GE Healthcare) at 25° C in HBS-EP+ buffer (10 mM HEPES pH 7.4, 150 mM NaCl, 3.4 mM EDTA, 0.005% (v/v) surfactant P20). For the measurement of human FcγR binding affinity, antibodies were immobilized on Series S Protein G sensor chip (GE Healthcare) at a density of 2000 RU. Serial two-fold dilutions of soluble, recombinant human FcγR ectodomains were injected as analytes. The FcγRs tested were FcγRI (10256-H08H, Sino Biological), FcγRIIAR131(10374-H08H1, Sino Biological), FcγRIIAH131 (10374-H08H1, Sino Biological), FcγRIIB (10259-H08H, Sino Biological), and FcγRIIIAV158 (10389-H08H1, Sino Biological) and FcγRIIIAF158 (10389-H08H, Sino Biological). The FcγRs were injected through flow cells at a flow rate of 20 μl min-1 with the concentration ranging from 0.488 – 250 nM (serial two-fold dilutions) for FcγRI, and 15.625 – 1000 nM (serial two-fold dilutions) for all the other FcγRs. A concentration ranging from 175.6 – 3000 nM (serial 1.5 fold dilutions) was used to test IgG2 binding affinity to FcγRIIIaF158 and FcγRIIIaV158. Association time was 60 s followed by 300 s dissociation (900 s for FcγRI). At the end of each cycle, sensor surface was regenerated with glycine HCl buffer (10 mM, pH 1.5) at a flow rate of 50 μl min-1 for 30 s. Background binding to blank immobilized flow cells was subtracted and affinity constants were calculated using BIA Evaluation software (GE Healthcare) using the Steady State Affinity model. For the measurement of human FcRn binding affinity, antibodies were immobilized on Series S Protein L sensor chip (GE Healthcare) at a density of 400 RU and experiments were performed in HBS-EP+ buffer at pH 6.0 following the method described above. Human FcRn/β2m (Sino Biological) was tested at a concentration ranging from 17.5 – 200 nM (serial 1.5-fold dilutions).

### In vivo cellular depletion assays

The *in vivo* cytotoxic activity of Fc glycovariants of mAbs was assessed in either FcγR humanized or FcRn/FcγR humanized mice in models of mAb-mediated depletion of CD4+ T cells or platelets, following previously described protocols (SMITH 2012 PNAS). For the T cell depletion model, mice were injected i.p. with 50 μg of recombinant GK1.5 human IgG1 mAb Fc variants and the abundance of CD4+ CD8− T cells were determined 48 h post-mAb administration in the blood and spleen by flow cytometric analysis. Baseline CD4+ CD8− T cell frequency was determined in blood samples obtained prior to mAb administration. For the platelet depletion model, mice were injected i.v. with the indicated dose of recombinant 6A6 human IgG1 mAb Fc variants. Mice were bled at the indicated timepoints before and after mAb administration, and platelet counts were measured using an automated hematologic analyzer (Advia 120 system or Heska HT5).

### Cloning, expression, and purification of recombinant IgG antibodies

Recombinant antibodies were generated following previously described protocols (34). Briefly, antibodies were generated by transient transfection of Expi293 cells with heavy and light chain expression plasmids. Prior to transfection, plasmid sequences were validated by direct sequencing (Genewiz). Recombinant IgG antibodies were purified from cell-free supernatants by affinity purification using Protein G or Protein A sepharose beads (GE Healthcare). Purified proteins were dialyzed in PBS, filter-sterilized (0.22 μm), and purity was assessed by SDS-PAGE followed by Coomasie blue staining. All antibody preparations were >90% pure and endotoxin levels were <0.005 EU/mg, as determined by the Limulus Amebocyte Lysate (LAL) assay.

### Quantification of serum and tissue IgG levels

Blood from mice was collected into gel microvette tubes and serum was fractionated by centrifugation (10,000*g*, 5 min). Fetal tissue lysates were prepared by mechanical homogenization in ice cold lysis buffer (50 mM Tris, 150 mM NaCl, 1% Triton X-100, 1 % Y-30, pH 7.4) using gentleMACS M tubes. Lysates were clarified by centrifugation (10,000*g*, 10 min) and supernatants were stored at −20°C. IgG levels in serum samples and tissue homogenates were determined by ELISA following previously published protocols (35). Briefly, high-binding 96-well microtiter plates (Nunc) were coated overnight at 4°C with Neutravidin (2 μg/ml in PBS).

All sequential steps were performed at room temperature. Plates were blocked for 1 h with PBS/2% BSA and incubated with biotinylated goat anti-human IgG antibodies for 1 h (5 μg/ml). Serum or tissue lysate samples were serially diluted and incubated for 1 h, followed by incubation with horseradish peroxidase-conjugated anti-human IgG (1:5000). Plates were developed using the TMB (3,3’,5,5’-Tetramethylbenzidine) two-component peroxidase substrate kit (KPL) and reactions stopped with the addition of 1 M phosphoric acid. Absorbance at 450nm was immediately recorded using a SpectraMax Plus spectrophotometer (Molecular Devices) and background absorbance from negative control samples was subtracted. Detection was performed using TMB (3,3’,5,5’-Tetramethylbenzidine) two-component peroxidase substrate kit (KPL) and reactions stopped with the addition of 2M phosphoric acid. Absorbance at 450nm was immediately recorded using a SpectraMax Plus spectrophotometer (Molecular Devices), background absorbance from negative control samples was subtracted and duplicate wells were averaged.

### Statistical Analysis

Results from multiple experiments are presented as mean ± standard error of the mean (SEM). One- or two-way ANOVA was used to test for differences in the mean values of quantitative variables, and where statistically significant effects were found, post-hoc analysis using Bonferroni multiple comparison test was performed. Data were analyzed with Graphpad Prism software (Graphpad) and P values of <0.05 were considered to be statistically significant.

## Acknowledgments

We would like to thank P. Smith, E. Lam, R. Peraza, and M. Ye (Rockefeller University) for excellent technical help. We acknowledge support from the Rockefeller University, Stanford University and the Chan Zuckerberg Biohub. We thank Dr. Grant Dorsey (UCSF), Prof Moses Kamya, the Infectious Diseases Research Collaboration, and the PROMOTE study team for provision of paired maternal and cord blood clinical samples. Pf CSP protein was provided by Drs. David Narum and Patrick Duffy at the National Institutes of Health, Laboratory of Malaria Immunology and Vaccinology. We thank members of the study team based at the Hospital de la Mujer Bertha Calderon Roque; the Hospital Infantil Manuel de Jesús Rivera; the Sócrates Flores Vivas, Edgar Lang, Francisco Buitrago, and Via Libertad Health Centers; the National Virology Laboratory in the Centro Nacional de Diagnóstico y Referencia, and the Sustainable Sciences Institute in Nicaragua for their dedication and high-quality work, as well as the Nicaraguan Ministry of Health and especially the study participants. Research reported in this publication was supported by the Bill and Melinda Gates Foundation grant OPP1188461 (to J.V.R. and T.T.W.) and in part by the National Institute of Allergy and Infectious Diseases (R01AI129795 to J.V.R; R01AI139119 to T.T.W; U19AI118610 to E.H.) and the National Institute of General Medical Science (R01GM096973 to L.W.). The content is solely the responsibility of the authors and does not necessarily represent the official views of the NIH.

## Author Contributions

S. Borghi, S. Bournazos, T.T.W., and J.V.R. designed experiments. S. Borghi, S. Bournazos, N.K.T., C.L., R.S., S.Z., L.W., T.T.W., and J.V.R. collected and/or analyzed data. C.L., A.G., E.H., and P.J., provided essential reagents. S. Borghi, S. Bournazos, T.T.W., and J.V.R wrote the manuscript.

---

**Table S1.** Characteristics of patient cohorts Table related to Figures 1–3.

| Paired Maternal/<br>Cord Subject ID | Gestational age at delivery in weeks | Maternal age at delivery in years | Gravidity | Maternal malaria infection during pregnancy | Gestational age at maternal infection (measured at every 4 week visits) | Placental malaria by histopathology | Baby sex |
| --- | --- | --- | --- | --- | --- | --- | --- |
| 1 | 38 | 22.6 | 3 | Malaria + | 16 weeks | - | F |
| 3 | 39 | 17.4 | 1 | Malaria + | 20 weeks | + | M |
| 9 | 39 | 28.3 | 2 | Malaria + | 24-36 weeks | + | M |
| 10 | 39 | 16.4 | 1 | Malaria - | None | - | M |
| 19 | 39 | 17.6 | 1 | Malaria + | 16 weeks | + | F |
| 36 | 36 | 17.7 | 1 | Malaria + | 20-24 weeks | + | F |
| 50 | 32 | 21.5 | 2 | Malaria - | None | - | M |
| 52 | 39 | 18.4 | 1 | Malaria + | 16-28 weeks | + | F |
| 58 | 39 | 17.7 | 1 | Malaria + | 20-28 weeks | + | M |
| 98 | 33 | 19.7 | 1 | Malaria + | 20 weeks | + | M |
| 106 | 38 | 20.5 | 2 | Malaria + | 16-36 weeks | + | M |
| 160 | 33 | 17.0 | 1 | Malaria + | 16 weeks | - | M |

**B**
| Code for maternal/<br>cord paired samples | Maternal/cord paired sample set | Gestational age at delivery | Maternal age at delivery | First pregnancy | Gravidity | Gestational age at maternal illness | Baby sex |
| --- | --- | --- | --- | --- | --- | --- | --- |
| ZP001 | 1 | 40.6 | 39 | No |  | 21 weeks 6 days | F |
| ZP005 | 2 | 38.1 | 18 | No |  | 14 weeks | F |
| ZP007 | 3 | 38.3 | 24 | No |  | 8 weeks | M |
| ZP008 | 4 | 39.3 | 19 | Yes |  | 19 weeks | M |
| ZP009 | 5 | 37.2 | 22 | No |  | 15 weeks | M |
| ZP010 | 6 | 38.2 | 18 | Yes |  | 12 weeks | F |
| ZP011 | 7 | 38.6 | 22 | No |  | 9 weeks | F |
| ZP017 | 8 | 38.1 | 33 | No |  | 4 weeks 2 days | F |
| ZP019 | 9 | 39.5 | 28 | No |  | 13 weeks | F |
| ZP021 | 10 | 39.3 | 25 | Yes |  | 8 weeks | F |
| ZP022 | 11 | 39.5 | 30 | Yes |  | 11 weeks | F |
| ZP032 | 12 | 39.3 | 27 | No |  | 19 weeks 6 day | F |
| ZP041 | 13 | 41.2 | 19 | Yes |  | 16 weeks | M |
| ZP042 | 14 | 37.1 | 38 | No |  | 9 weeks | F |
| ZP043 | 15 | 39.0 | 27 | No |  | 2 weeks 4 days | M |
| ZP044 | 16 | 37.0 | 25 | Yes |  | 1 week 6 days | F |

**Table S2.**
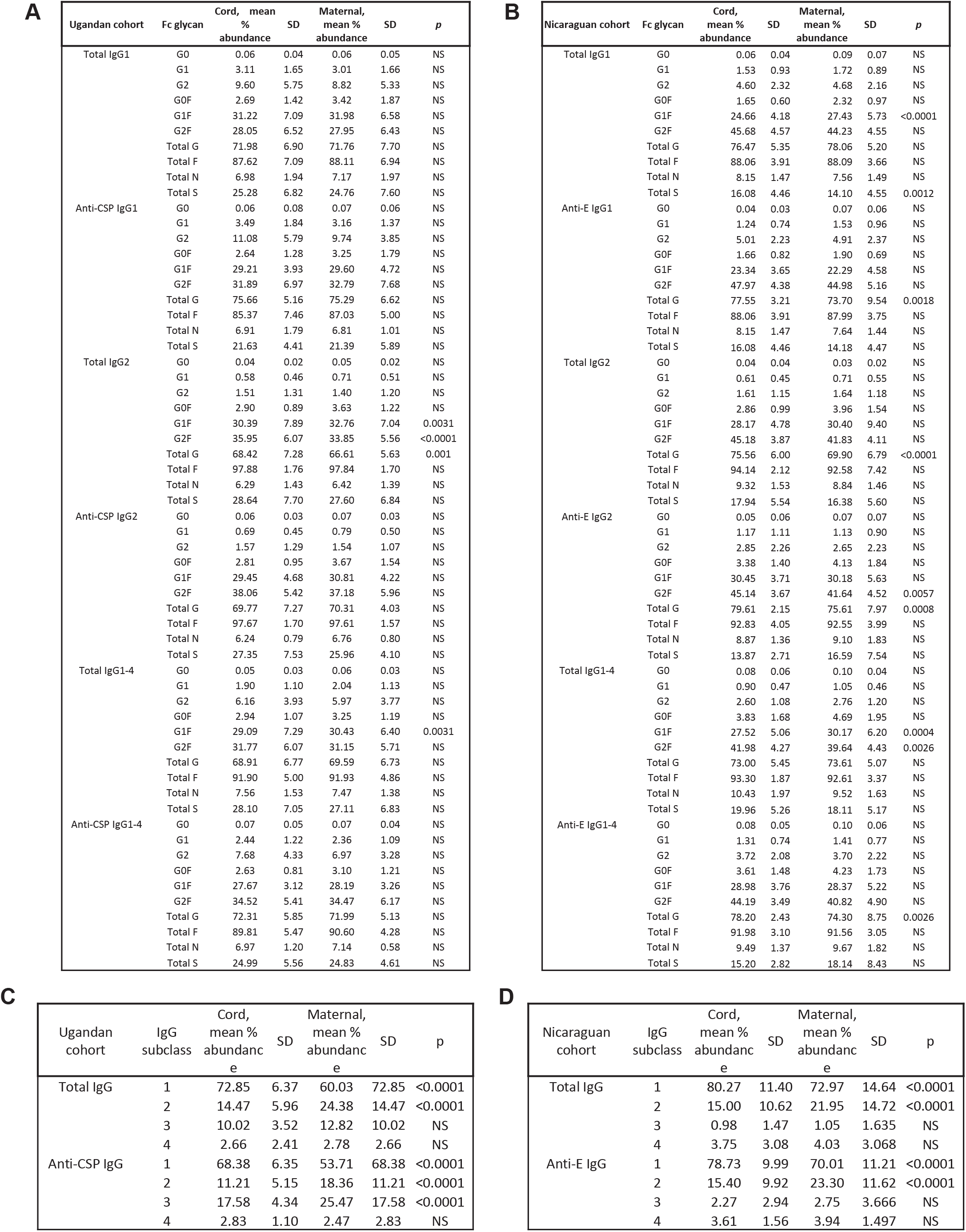
Quantification of paired maternal and cord IgG Fc glycans and IgG subclasses from two clinical cohorts. **(A-B)** All paired maternal and cord bloods were characterized for Fc glycosylation of total IgG1, IgG2 and IgG1-4 as well as **(A – Ugandan cohort)** anti-CSP IgG1, anti-CSP IgG2, and anti-CSP IgG1-4 or **(B – Nicaraguan cohort)** anti-Zika virus envelope (E) IgG1, anti-E IgG2, and anti-E IgG1-4. The relative abundance of the following Fc glycoforms were characterized for each IgG category: G0, G1, G2, G0F, G1F, G2F, total galactosylation (G), total fucosylation (F), total bisection (N), total sialylation (S). Significance was assessed by 2-way ANOVA with Bonferroni’s multiple comparisons test where p<0.0125 was considered significant, *p < 0.0125, **p < 0.0025, ***p < 0.00025. (**C-D**) Quantification of IgG subclasses in paired maternal and cord bloods. (**C**) Total or anti-malaria circumsporozoite protein (CSP) IgG from the Ugandan cohort. (**D**) Total or anti-Zika virus envelope (E) IgG from the Nicaraguan cohort. Significance was assessed by paired T-test with Bonferroni’s correction. \**P* < 0.05, \*\**P* < 0.01, \*\*\**P* < 0.001, \*\*\*\**P* < 0.0001. Table related to Figures 1–3 and Table S1.

**Figure S1:**
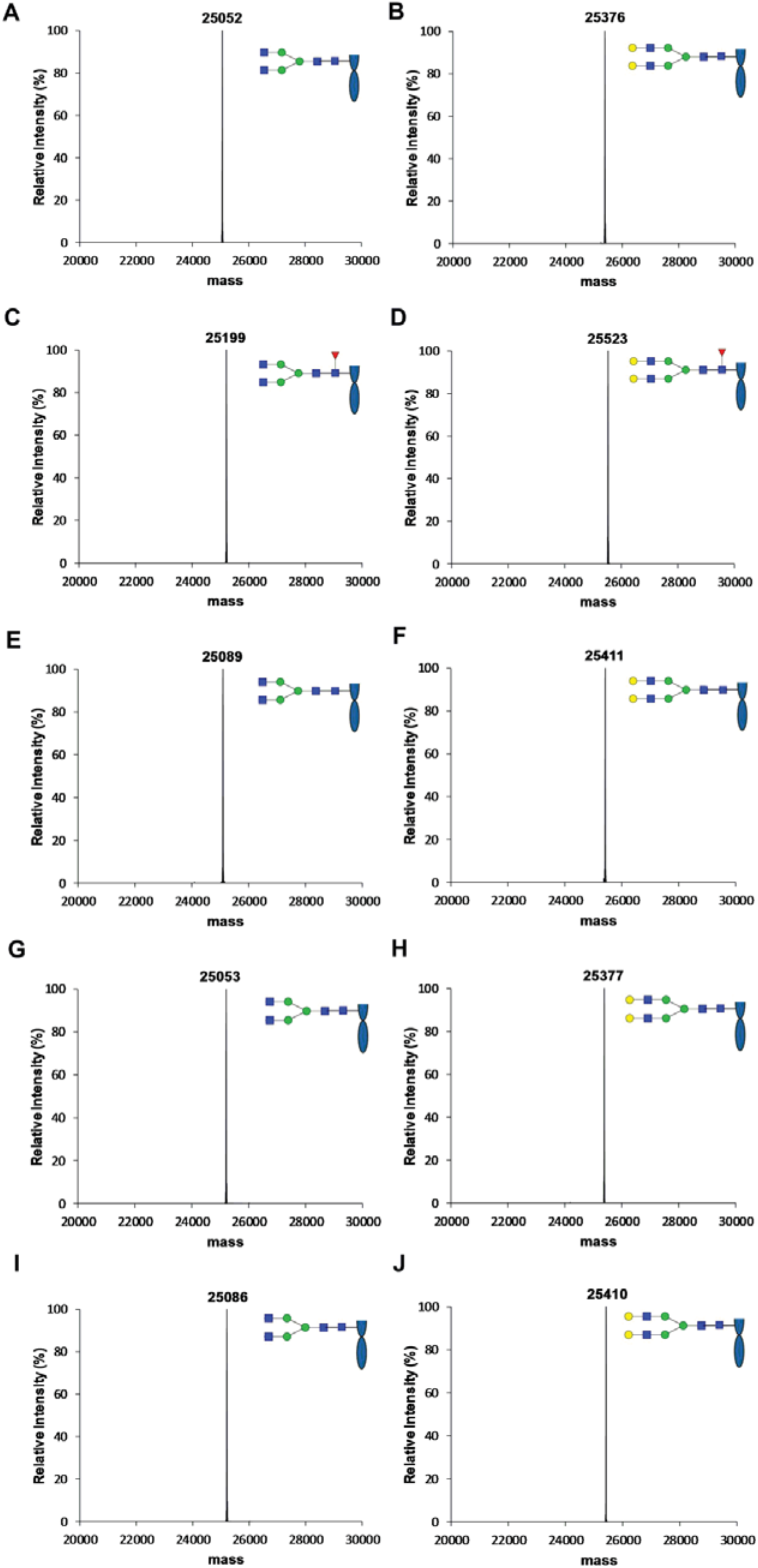
LC-ESI-MS analysis of the Fc domain from the glycoforms of the engineered 6A6 and FI6 after IdeS treatment. (**A**) 6A6_IgG1_G0. (**B**) 6A6_IgG1_G2. (**C**) 6A6_IgG1_G0F. (**D**) 6A6_IgG1_G2F. (**E**) 6A6_IgG2_G0. (**F**) 6A6_IgG2_G2. (**G**) FI6_IgG1_G0. (**H**) FI6_IgG1_G2. (**I**) FI6_IgG2_G0. (**J**) FI6_IgG2_G2. The deconvoluted spectra of the Fc fragments are shown. Figure related to Figures 4–6.

**Figure S2:**
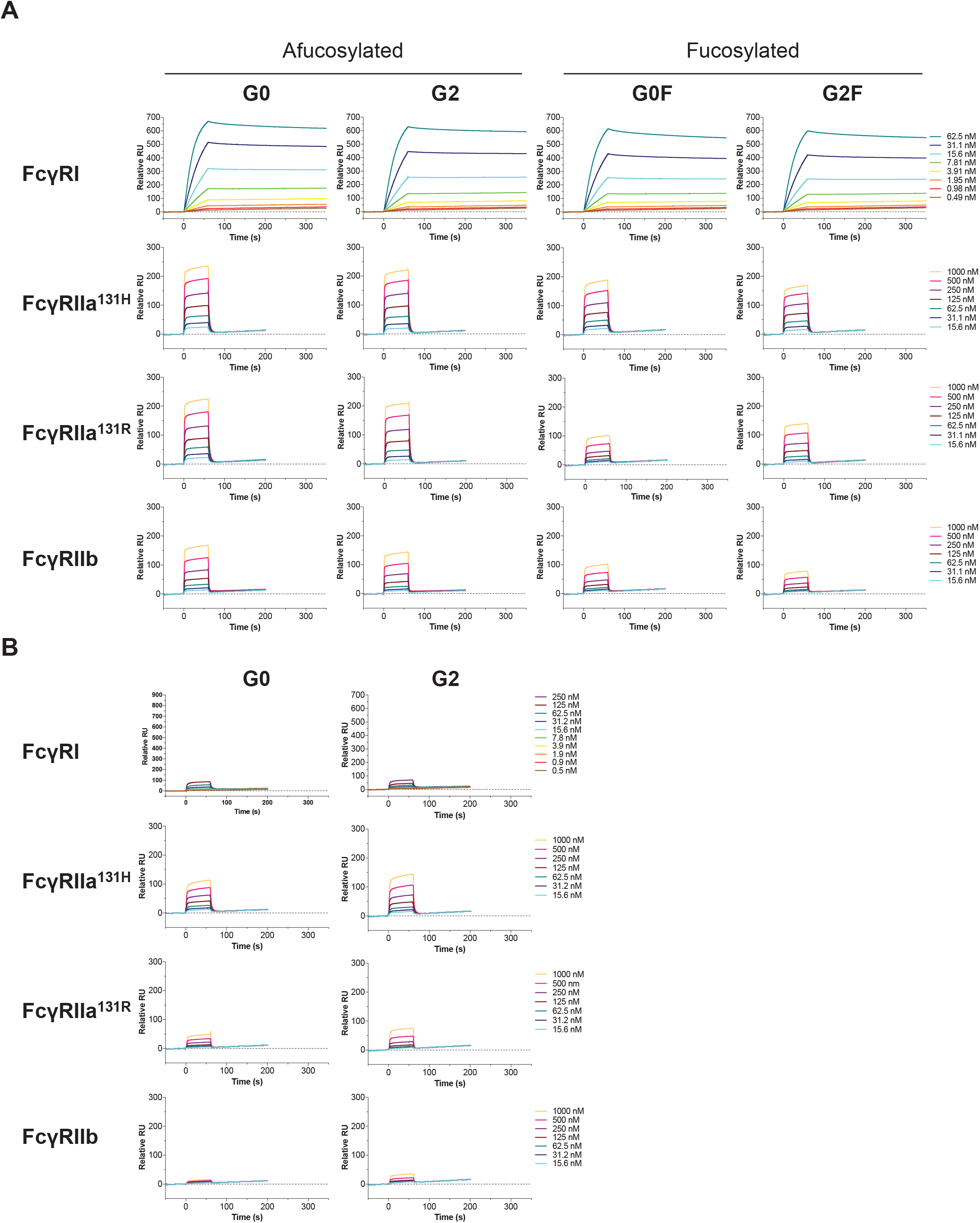
Effect of IgG1 and IgG2 core fucosylation and galactosylation on FcγRs engagement. The affinity of glycoengineered mouse-human chimeric IgG1 (**A**) and IgG2 (**B**) of anti-gpIIb (6A6) mAb for human FcγRI, FcγRIIa^131H^, FcγRIIa^131R^ and FcγRIIb ectodomains was determined by SPR analysis and representative SPR sensorgrams are presented. Figure related to Figures 4 and 5.

**Figure S3:**
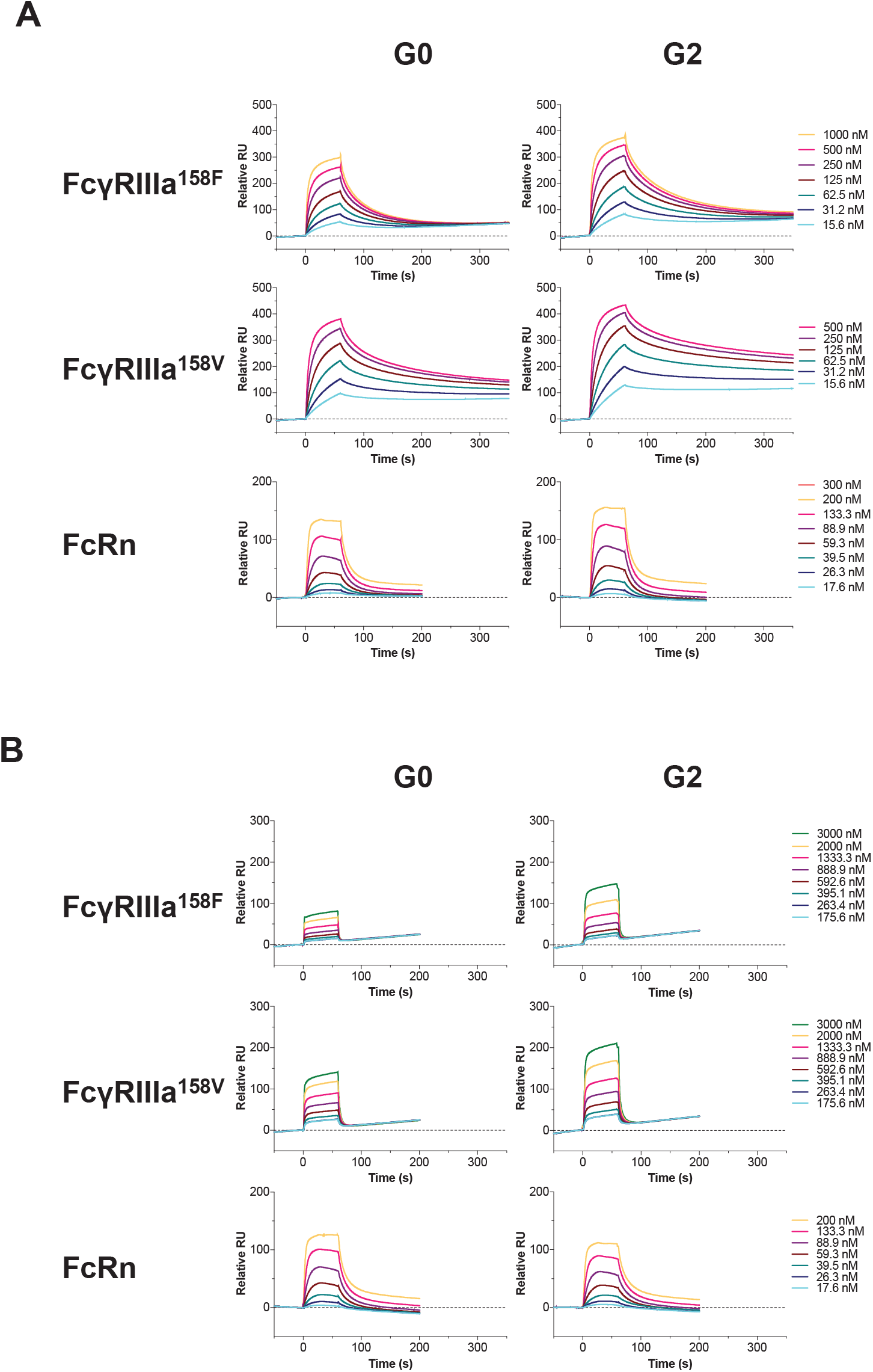
Effect of IgG core fucosylation and galactosylation on FcγRs engagement. The affinity of glycoengineered human IgG1 (**A**) and IgG2 (**B**) of anti-HA FI6 mAb for human FcγRIIIa^158F^, FcγRIIa^158V^ and FcRn ectodomains was determined by SPR analysis and representative SPR sensorgrams are presented. Figure related to Figures 4 and 5.

**Figure S4:**
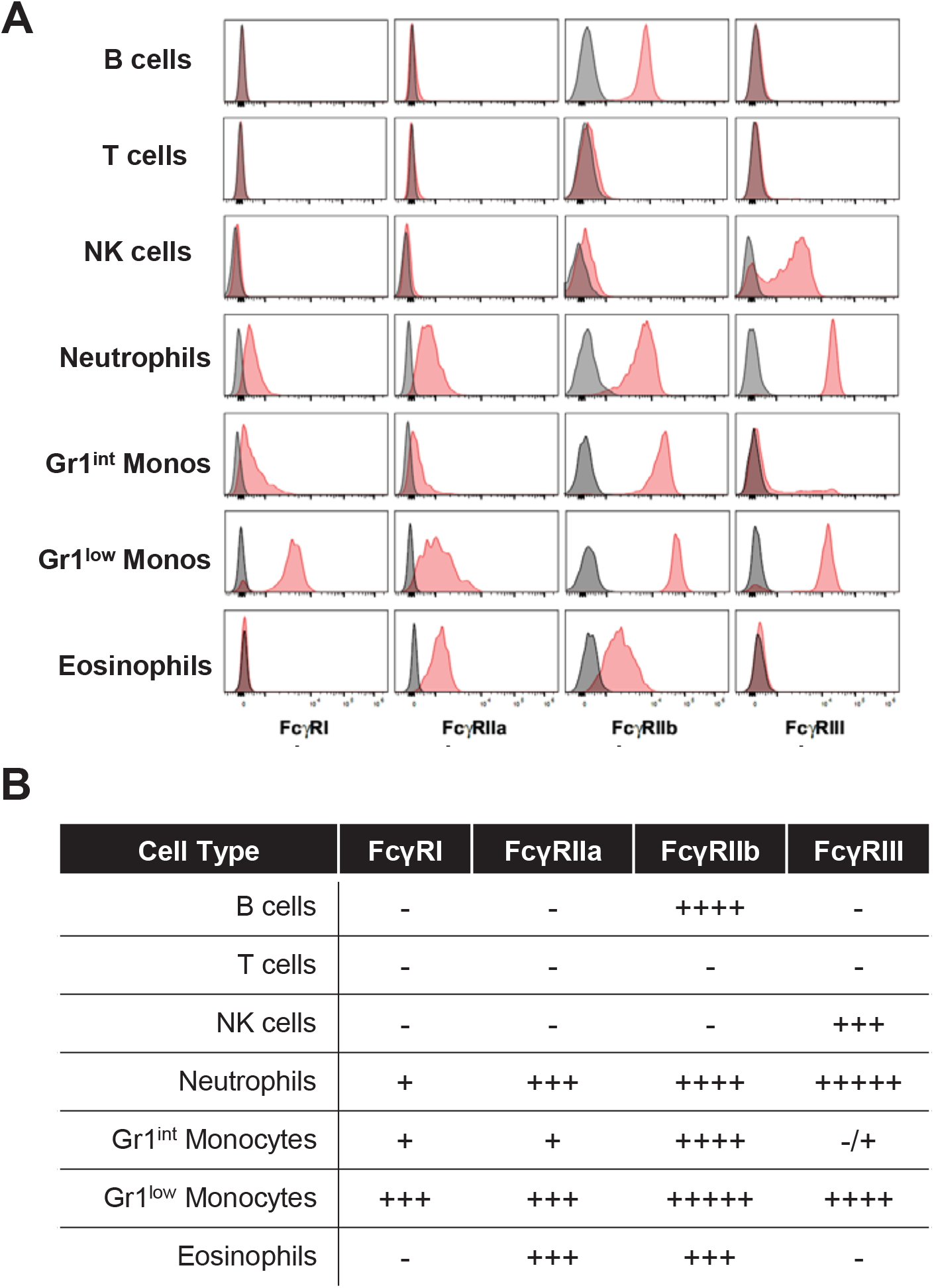
Characterization of the FcγR expression profile of FcRn/FcγR humanized mice. Leukocyte populations from FcRn/FcγR humanized mice were characterized for their expression of the various human FcγRs by flow cytometry. (**A**) Representative flow cytometry overlay histograms of human FcγR expression (black: isotype control; red: anti-human FcγR mAbs) in different circulating leukocyte populations (B cells (CD3^-^/CD11b^-^/B220^+^); T cells (CD3^+^/B220^-^ /CD11b^-^); NK cells (CD3^-^/B220^-^/NK1.1^+^); Neutrophils (CD3^-^/B220^-^/NK1.1^-^ /CD11b^+^/Gr1^high^/SSC^high^); Gr1^int^ monocytes (CD3^-^/B220^-^/NK1.1^-^/CD11b^+^/Gr1^int^/SSC^low^); Gr1^low^ monocytes (CD3^-^/B220^-^/NK1.1^-^/CD11b^+^/Gr1^low^/SSC^low^); Eosinophils (CD3^-^B220^-^/NK1.1^-^ /CD11b^+^/Gr1^low^/SSC^high^)). (**B**) Overview of the human FcγR expression pattern of leukocyte populations from FcRn/FcγR humanized mice. Figure related to Figures 6 and 7.

**Figure S5:**
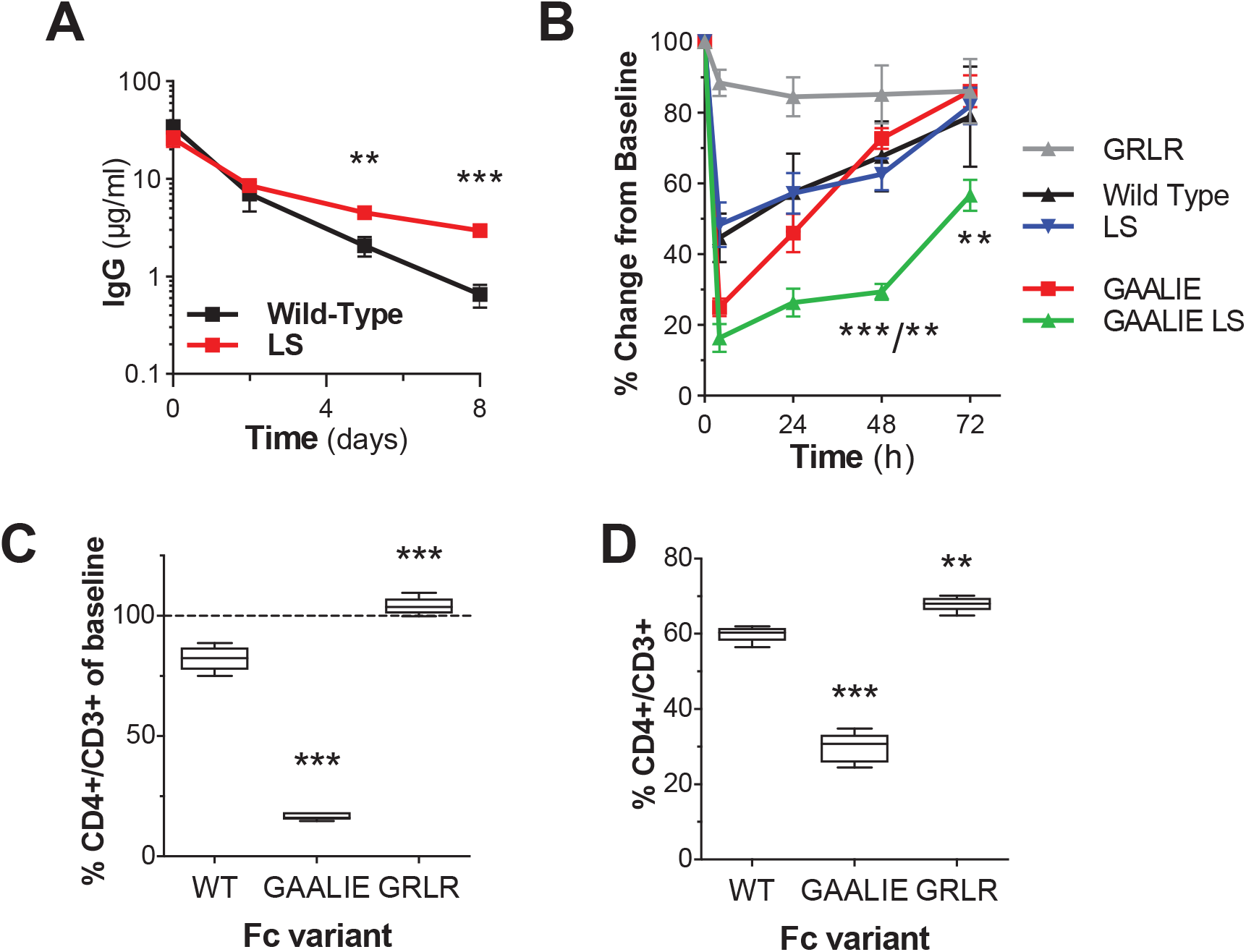
Evaluation of the *in vivo* half-life and Fc effector function of human IgG antibodies in FcRn/FcγR humanized mice. The *in vivo* function of Fc engineered human IgG1 antibodies was evaluated in FcRn/FcγR humanized mice to confirm the functional activity of human FcRn and FcγRs. (**A**) Fc domain variants (LS: M428L/N434S) with increased affinity for human FcRn were generated for the human IgG1 anti-HIV mAb 3BNC117 and their *in vivo* halflife was evaluated in FcRn/FcγR humanized mice. Consistent with its increased affinity for human FcRn, the LS variant exhibited extended half-life in FcRn/FcγR humanized mice. Results are presented as the mean ± SEM from 5-6 mice/group. ns= not significant; *** p<0.01; ****p=0.0002. (**B-D**) The cytotoxic activity of Fc engineered variants with differential FcγR and FcRn binding affinity was evaluated in FcRn/FcγR humanized mice in models of mAb-mediated cytotoxic depletion of mouse platelets (**B**) and CD4^+^ T cells (**C**-**D**). (**B**) Timecourse analysis of platelet counts of FcRn/FcγR humanized mice following administration of Fc engineered anti-platelet mAbs with differential FcγR and FcRn affinity. Results are presented as the mean ± SEM from 5 mice/group; *** p<0.0001 vs. GAALIE; ** p=0.006 vs. LS; *p=0.01 vs. GAALIE.CD4^+^ T cell counts were determined in the blood (**C**) and spleen (**D**) of FcRn/FcγR humanized mice 24 h following administration of Fc domain variants of anti-CD4 mAb (WT: baseline FcγR affinity; GRLR: diminished FcγR affinity; GAALIE: enhanced FcγR affinity). 5 mice/group; **** p<0.0001; ** p=0.002. Figure related to Figures 6 and 7.

## References

1. V. Ghetie, E. S. Ward, FcRn: the MHC class I-related receptor that is more than an IgG transporter. Immunol Today 18, 592–598 (1997).

2. N. E. Simister, E. Jacobowitz Israel, J. C. Ahouse, C. M. Story, New functions of the MHC class I-related Fc receptor, FcRn. Biochem Soc Trans 25, 481–486 (1997).

3. A. P. West, Jr., P. J. Bjorkman, Crystal structure and immunoglobulin G binding properties of the human major histocompatibility complex-related Fc receptor(,). Biochemistry 39, 9698–9708 (2000).

4. T. Bakchoul et al., Inhibition of HPA-1a alloantibody-mediated platelet destruction by a deglycosylated anti-HPA-1a monoclonal antibody in mice: toward targeted treatment of fetal-alloimmune thrombocytopenia. Blood 122, 321–327 (2013).

5. L. C. Simmons et al., Expression of full-length immunoglobulins in Escherichia coli: rapid and efficient production of aglycosylated antibodies. Journal of immunological methods 263, 133–147 (2002).

6. M. H. Tao, S. L. Morrison, Studies of aglycosylated chimeric mouse-human IgG. Role of carbohydrate in the structure and effector functions mediated by the human IgG constant region. Journal of immunology 143, 2595–2601 (1989).

7. S. Bournazos, T. T. Wang, R. Dahan, J. Maamary, J. V. Ravetch, Signaling by Antibodies: Recent Progress. Annual review of immunology 35, 285–311 (2017).

8. T. T. Wang, IgG Fc Glycosylation in Human Immunity. Current topics in microbiology and immunology (2019).

9. H. K. Einarsdottir et al., Comparison of the Fc glycosylation of fetal and maternal immunoglobulin G. Glycoconj J 30, 147–157 (2013).

10. C. Fu et al., Placental antibody transfer efficiency and maternal levels: specific for measles, coxsackievirus A16, enterovirus 71, poliomyelitis I-III and HIV-1 antibodies. Sci Rep 6, 38874 (2016).

11. V. Watanaveeradej et al., Transplacentally transferred maternal-infant antibodies to dengue virus. The American journal of tropical medicine and hygiene 69, 123–128 (2003).

12. J. P. van den Berg et al., Transplacental transport of IgG antibodies specific for pertussis, diphtheria, tetanus, haemophilus influenzae type b, and Neisseria meningitidis serogroup C is lower in preterm compared with term infants. Pediatr Infect Dis J 29, 801–805 (2010).

13. S. C. Stach et al., Placental transfer of IgG antibodies specific to Klebsiella and Pseudomonas LPS and to group B Streptococcus in twin pregnancies. Scandinavian journal of immunology 81, 135–141 (2015).

14. M. F. Jennewein et al., Fc Glycan-Mediated Regulation of Placental Antibody Transfer. Cell 178, 202–215 e214 (2019).

15. D. R. Martinez et al., Fc Characteristics Mediate Selective Placental Transfer of IgG in HIV-Infected Women. Cell 178, 190–201 e111 (2019).

16. A. Kakuru et al., Dihydroartemisinin-Piperaquine for the Prevention of Malaria in Pregnancy. The New England journal of medicine 374, 928–939 (2016).

17. P. Jagannathan et al., Dihydroartemisinin-piperaquine for intermittent preventive treatment of malaria during pregnancy and risk of malaria in early childhood: A randomized controlled trial. PLoS Med 15, e1002606 (2018).

18. A. Balmaseda et al., Antibody-based assay discriminates Zika virus infection from other flaviviruses. Proc Natl Acad Sci U S A 114, 8384–8389 (2017).

19. T. T. Wang et al., Anti-HA Glycoforms Drive B Cell Affinity Selection and Determine Influenza Vaccine Efficacy. Cell 162, 160–169 (2015).

20. P. Palmeira, C. Quinello, A. L. Silveira-Lessa, C. A. Zago, M. Carneiro-Sampaio, IgG placental transfer in healthy and pathological pregnancies. Clin Dev Immunol 2012, 985646 (2012).

21. H. K. Einarsdottir et al., On the perplexingly low rate of transport of IgG2 across the human placenta. PloS one 9, e108319 (2014).

22. S. Hashira, S. Okitsu-Negishi, K. Yoshino, Placental transfer of IgG subclasses in a Japanese population. Pediatr Int 42, 337–342 (2000).

23. R. L. Shields et al., Lack of fucose on human IgG1 N-linked oligosaccharide improves binding to human Fcgamma RIII and antibody-dependent cellular toxicity. The Journal of biological chemistry 277, 26733–26740 (2002).

24. T. Shinkawa et al., The absence of fucose but not the presence of galactose or bisecting N-acetylglucosamine of human IgG1 complex-type oligosaccharides shows the critical role of enhancing antibody-dependent cellular cytotoxicity. The Journal of biological chemistry 278, 3466–3473 (2003).

25. G. Dekkers et al., Affinity of human IgG subclasses to mouse Fc gamma receptors. MAbs 9, 767–773 (2017).

26. T. Li et al., Modulating IgG effector function by Fc glycan engineering. Proc Natl Acad Sci U S A 114, 3485–3490 (2017).

27. C. Li, L. X. Wang, Chemoenzymatic Methods for the Synthesis of Glycoproteins. Chem Rev 118, 8359–8413 (2018).

28. M. Olsson, P. Bruhns, W. A. Frazier, J. V. Ravetch, P. A. Oldenborg, Platelet homeostasis is regulated by platelet expression of CD47 under normal conditions and in passive immune thrombocytopenia. Blood 105, 3577–3582 (2005).

29. P. Smith, D. J. DiLillo, S. Bournazos, F. Li, J. V. Ravetch, Mouse model recapitulating human Fcgamma receptor structural and functional diversity. Proceedings of the National Academy of Sciences of the United States of America 109, 6181–6186 (2012).

30. I. Schwab, A. Lux, F. Nimmerjahn, Pathways Responsible for Human Autoantibody and Therapeutic Intravenous IgG Activity in Humanized Mice. Cell Rep 13, 610–620 (2015).

31. J. Zalevsky et al., Enhanced antibody half-life improves in vivo activity. Nature biotechnology 28, 157–159 (2010).

32. S. Bournazos et al., Broadly neutralizing anti-HIV-1 antibodies require Fc effector functions for in vivo activity. Cell 158, 1243–1253 (2014).

33. T. T. Wang et al., IgG antibodies to dengue enhanced for FcgammaRIIIA binding determine disease severity. Science 355, 395–398 (2017).

34. S. Bournazos, D. J. DiLillo, A. J. Goff, P. J. Glass, J. V. Ravetch, Differential requirements for FcγR engagement by protective antibodies against Ebola virus. Proc Natl Acad Sci U S A 116, 20054–20062 (2019).

35. P. Weitzenfeld, S. Bournazos, J. V. Ravetch, Antibodies targeting sialyl Lewis A mediate tumor clearance through distinct effector pathways. J Clin Invest 129, 3952–3962 (2019).

